# Inferring toxicant susceptibility in global populations from gene-environment interactions involving the Aryl Hydrocarbon Receptor

**DOI:** 10.64898/2026.01.10.698796

**Authors:** Uchechukwu S. Chimeh, Molly C. Rogers, David L. Aylor

## Abstract

Gene-environment interaction (GxE) studies comprise a very small part of the genetics or environmental epidemiology literature, and most existing studies are in populations of European ancestry. We made predictions about GxE in global populations by combining existing GxE studies with genetic data from the 1000 Genomes Project (1kGP), which captured genetic variation in diverse populations worldwide. We modeled susceptibility of 1kGP populations to 2, 3, 7, 8-tetrachlorodibenzo-p-dioxin (TCDD) exposures based on variation in the aryl hydrocarbon receptor (*AHR*) gene. The premise of our approach is that the risk variants involved in GxE are shared across global super-populations but vary in frequency by population. We built our model upon GxE estimates from a study in Seveso, Italy, where offspring birthweight was influenced by *AHR* variants after TCDD exposure. Our simulations predicted that GxE would result in different outcomes across global populations. This framework can be extended to model population susceptibility to a broad range of toxicants that impact public health, including common AHR ligands like the polycyclic aromatic hydrocarbons found in cigarette smoke and diesel exhaust.

## Introduction

Genetic diversity influences traits including our appearances, predispositions to diseases, and susceptibilities to the adverse effects of environmental exposures. However, most genetic associations have been mapped in populations of European descent (1), leaving knowledge gaps concerning global populations. The human genome-wide association study (GWAS) catalog, which houses results from studies associating genetic variants with traits, has a population breakdown of 79% European, 8% East Asian, 2% African, 2% South Asian, and 1% Hispanic or Latin American individuals (2, 3). Due to the skewed representation of global populations in genetic association studies, it is challenging to fully understand how genetic risk variants contribute to trait variation and health disparities.

One strategy is to model risk allele frequencies across diverse populations. Most common genetic variants are shared among human populations and population reference panels have established how the frequencies of these variants differ globally. One such reference panel is the 1000 Genomes Project (1kGP). The 1kGP is the largest public repository of human genetic variation data. It widely sampled various global populations to identify genetic variants with frequencies of at least 1%. Most common variants within the human genome are shared between the five super-populations that comprise the 1kGP (African, Ad-mixed American, East Asian, European, and South Asian) (4). These shared variants offer critical insights into risk and susceptibility in understudied populations. Our premise is that higher frequencies of risk variants will correspond to higher public health impact on populations. Inferring genetic effects across populations is especially relevant to gene-environment interaction (GxE) studies, which focus on how genetic variants mediate response to environmental exposures. Higher frequencies of risk variants at critical environmental response genes could increase the incidence of adverse outcomes after exposure to toxicants. GxE studies are a small subset of association studies (5), and the paucity of data in most populations motivated us to extend existing results to new populations.

We extended existing GxE results from the Seveso Women’s Health Study (SWHS) to account for variance in risk allele frequencies among global populations. The SWHS followed the population in and around Seveso, Italy that was exposed to high levels of 2, 3, 7, 8-tetrachlorodibenzo-p-dioxin (TCDD) after a factory explosion in 1976 (6). A candidate gene association study based on SWHS data discovered that six single nucleotide polymorphisms (SNPs) in the aryl hydrocarbon receptor (*AHR*) gene (rs6968865, rs3757824, rs10249788, rs2282885, rs2040623, and rs2106728) were risk variants and interacted with TCDD to decrease offspring birthweight (7). AhR is an important receptor in the metabolism of various xenobiotic compounds including TCDD, polycyclic aromatic hydrocarbons (PAHs), and others. *AHR* ligands bind to the inactive protein in the cytoplasm, which is then translocated to the nucleus to dimerize with the AhR nuclear translocator (*ARNT*) (8). The activated AHR-ARNT complex binds xenobiotic response elements in target genes like *CYP1A1* to regulate their expression (9). Many common environmental factors such as diesel exhaust, cigarette smoke, and waste from industrial burning processes contain *AHR* ligands, and it is highly relevant to global public health. Multiple genetic variants in the *AHR* gene have previously been associated with GxE effects on a range of traits (7, 10–12). A separate SWHS study found that the SNP rs6968865 decreased fertility and fecundity in TCDD exposed women (11). A study of Chinese coke oven workers linked rs2282885 to differences in urinary 1-hydroxypyrene after PAH exposure (10), which is a biomarker of internal dose. Here, we modeled birthweight decreases after simulated TCDD exposure in five 1kGP super-populations. We combined GxE estimates from the SWHS study (7) with *AHR* risk allele frequencies from 1kGP data, then randomly assigned TCDD dose to create simulated populations. We found that *AHR* allele frequencies varied widely and that GxE could cause differences in birthweight distributions between super-populations.

## Materials and Methods

### Collecting and processing *AHR* variant data from The 1000 Genomes Project

We retrieved genetic variants from a human population reference panel established by The 1000 Genomes Project (1kGP). Phase 3 of the 1kGP sequenced the genomes of 2504 individuals from 26 populations, grouped into five super-populations by continental origin: African (AFR), Ad-mixed American (AMR), East Asian (EAS), European (EUR), and South Asian (SAS). There were 661, 347, 504, 503 and 489 individuals respectively in the super-population groups. The 1kGP aligned sequence data to the GRCh37 human genome build and generated variant callsets from sequence data at a false discovery rate threshold of <5%. Callsets included biallelic and multi-allelic single nucleotide polymorphisms (SNPs), insertions and deletions (indels), multi-nucleotide polymorphisms (MNPs), complex substitutions and structural variants. These variants are available as phased haplotypes, which combine the variants found on the same chromosome, in variant call format (vcf) (4). 1kGP data have been used to analyze population genetic structure and to filter non-pathogenic variants from genetic data (13).

*AHR* is found on Chromosome 7 at the genomic coordinates of 16,916,359 and 17,346,152 in the GRCh38 human genome build, which maps to 16,955,983 to 17,385,776 in the GRCh37 build (14). We obtained Chromosome 7 variants and sample metadata from the 1kGP repository (4). We used vcftools (v0.1.16) (15) to filter biallelic *AHR* SNPs using the “--remove-indels” flag. We converted the variants to a PLINK transposed text genotype table (tped) and comma delimited format (csv). We used R (version 4.2.2) for downstream analyses.

### Calculating *AHR* nucleotide polymorphism, nucleotide diversity, and shared variation

We established how much variation exists globally in the *AHR* gene and how much of this variation was shared between 1000 Genomes super-populations. We estimated nucleotide polymorphism *P̂* by calculating the proportion of variant sites in the *AHR* gene. We then calculated the percentage of these variants shared between super-populations. To quantify how similar individuals were at the gene, we estimated nucleotide diversity [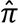] using the R package, pegas (v1.1) (16), correcting for the full length of *AHR*.

### Identifying functional *AHR* variants and their GxE effects from the Seveso Women’s Health Study

We focused on six *AHR* variants which interacted with TCDD to influence birthweight in the SWHS. Offspring birthweight decreased more if a mother harbored a risk genotype at *AHR* SNPs (rs6968865, rs3757824, rs10249788, rs2282885, rs2040623, and rs2106728), compared to offspring from mothers without the risk genotype who were exposed to similar levels of TCDD (7). We calculated the frequency of risk and non-risk alleles in the five 1000 Genomes super-populations. We identified each individual’s phased haplotypes, by combining SNPs on the same chromosome and compared the distribution of risk alleles and phased haplotypes from the six variants among the super-populations, using Europe as a proxy for the Seveso cohort. We determined how much of the haplotype pool was shared between Europe and other super-populations. This was done to ascertain if enough shared variation existed to extend the GxE effects on birthweight from Seveso to non-European populations.

Ames et al. used SWHS data to estimate the effect of *AHR*-TCDD interaction on birthweight using a dominant penetrance model in which individuals with at least one minor allele (homozygotes and heterozygotes) were compared to the major allele homozygotes (7). We used those GxE effect estimates to predict how birthweight would change in the super-populations if they were similarly exposed to TCDD. We calculated GxE effect per SNP (in grams/ log_10_ parts-per-trillion TCDD, g/log_10_ppt) as the difference in mean birthweight between individuals with the risk and non-risk genotypes for every unit of TCDD exposure (S1 Table). For example, rs6968865 mediated a 62.34g decrease in offspring birthweight per log_10_(TCDD exposure in ppt) from mothers with the risk genotypes (AA and AT). Offspring from mothers with the non-risk genotype (TT) had a non-significant increase in birthweight – the GxE model estimated an increase of 23.23g/log_10_(ppt) (7). We calculated this SNP’s GxE effect as the differences between the two estimates, -85.57 g/log_10_(ppt).

### Simulating TCDD exposure in the 1000 Genomes super-populations

We simulated TCDD exposure in the super-populations using summary statistics of serum TCDD concentration reported in Seveso. The SWHS collected serum samples archived by the Desio Hospital laboratory soon after the explosion. TCDD concentration ranged from 2.5 to 56000ppt with a 55.9ppt median in an approximately log-normal distribution (17). We simulated truncated log-normal TCDD distributions in the super-populations, using 1000 Genomes sample sizes and the log-transformed TCDD median and range. We used the rlnormTrunc function in R to create these distributions by calculating the mean and standard deviation of the log-transformed distributions from equations (1) and (2) respectively. We set a seed for reproducibility and iterated this model over 1000 simulations. Each subsequent seed was set as the next consecutive integer.

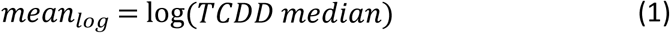

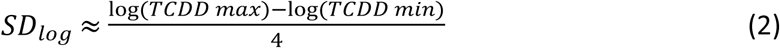

### Predicting GxE effect on birthweight in the 1000 Genomes super-populations

We modeled how birthweight would change among the super-populations after simulated TCDD exposure as mediated by each of the six functional *AHR* SNPs. To do this, we used the GxE effects estimated by the SWHS, the simulated TCDD exposure concentrations, risk genotype frequencies from the 1kGP, and simulated birthweight distributions.

First, we simulated birthweights for each individual in each super-population using random normal birthweight distributions with the SWHS mean and standard deviation (3264g ± 526g) (4, 7) and 1000 Genomes sample numbers. Using the R rnorm function, we set seeds and replicated 1000 birthweight distributions for each super-population. This yielded 5000 simulated populations evenly split between AFR, AMR, EAS, EUR, and SAS. Next, we set a seed and randomly sampled half the populations without replacement to model TCDD exposure. We simulated TCDD exposure for each individual in those 500 populations, yielding 500 simulations of exposed populations and 500 simulations of unexposed populations for each of the 5 super-populations.

Last, we calculated the effect of each SNP on birthweight using the random birthweight, random exposure level, and individual genotypes. The risk genotype of each SNP had a GxE effect (g/log_10_ppt) which varied by TCDD exposure as described above. Therefore, we multiplied SNP GxE effects by log_10_(TCDD) to get individual risk effects (in grams).

For each of the six super-populations, we tested for sampling variation by comparing birthweight means between random pairs of TCDD-unexposed and exposed simulated populations using student t-tests. We randomly drew one simulated unexposed population and one simulated exposed population representing the same super-population, and tested for a difference in birthweight means at each of six SNPs. Overall, we did 15,000 t-tests – 2500 tests for each GxE SNP evenly divided across the five super-populations. We summarized SNP effects by averaging the differences in birthweight means between unexposed and exposed populations in each of these tests. To test for differential effects across populations, we compared birthweight means across the five TCDD-exposed simulated super-populations using one-way ANOVAs. We randomly chose one simulated population representing each super-population for 500 tests per SNP. This resulted in 3000 ANOVA tests. To accommodate multiple tests, we adjusted the p-values for both sets of tests using the Benjamini-Hochberg false discovery rate (FDR) method and used a significance threshold of 0.05. All statistical tests were done using R (version 4.2.2).

## Results

### *AHR* sequence variation was abundant and was shared among 1000 Genomes super-populations

The *AHR* gene contained 13545 variants within its 429,793 nucleotides. These included 457 indels and 13008 SNPs, 13004 of which were biallelic. We computed nucleotide polymorphism, *P̂* = 0.0315. In other words, there were roughly three SNPs for every 100 bases in the *AHR* sequence on average. When we looked at this measure of genetic variation within each 1kGP super-population, we saw a range of values from 7.9×10^-3^ in East Asia to 1.5×10^-2^ in Africa (Table 1). Most SNPs occurred in multiple individuals (54.9%) (Fig 1a), and most of these non-singleton SNPs were shared across different populations (56.02%). A portion of these segregated in all five populations (14.33% of the total) (Fig 1b). These shared SNPs allow us to calculate nucleotide diversity (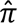), a population parameter that describes genetic similarity within a population. Nucleotide diversity for *AHR* in the 1kGP was 8.1×10^-4^, meaning that on average there was one difference every 1250 bases when all pairs of individuals were compared. Africa had the highest nucleotide diversity (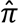 = 9.4×10-4), while East Asia had the least variation (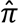 = 6.8×10-4) (Table 1). We concluded that *AHR* was highly polymorphic in human populations despite its functional importance. Furthermore, a substantial portion of these variants were shared among 1kGP super-populations, and those shared variants enabled our approach.

**Table 1.** Nucleotide polymorphism *P̂* and nucleotide diversity 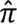 in the aryl hydrocarbon receptor gene in The 1000 Genomes Project (1kGP).

| Super-population | Sample Number | $\hat{P}$ ($\times 10^{-2}$) | $\hat{\pi}$ ($\times 10^{-4}$) |
| --- | --- | --- | --- |
| 1kGP | 2504 | 3.03 | 8.12 |
| African | 661 | 1.5 | 9.43 |
| Ad-mixed American | 347 | 0.99 | 7.08 |
| East Asian | 504 | 0.79 | 6.78 |
| European | 503 | 0.81 | 7.03 |
| South Asian | 489 | 0.9 | 7.35 |

**Fig 1.**
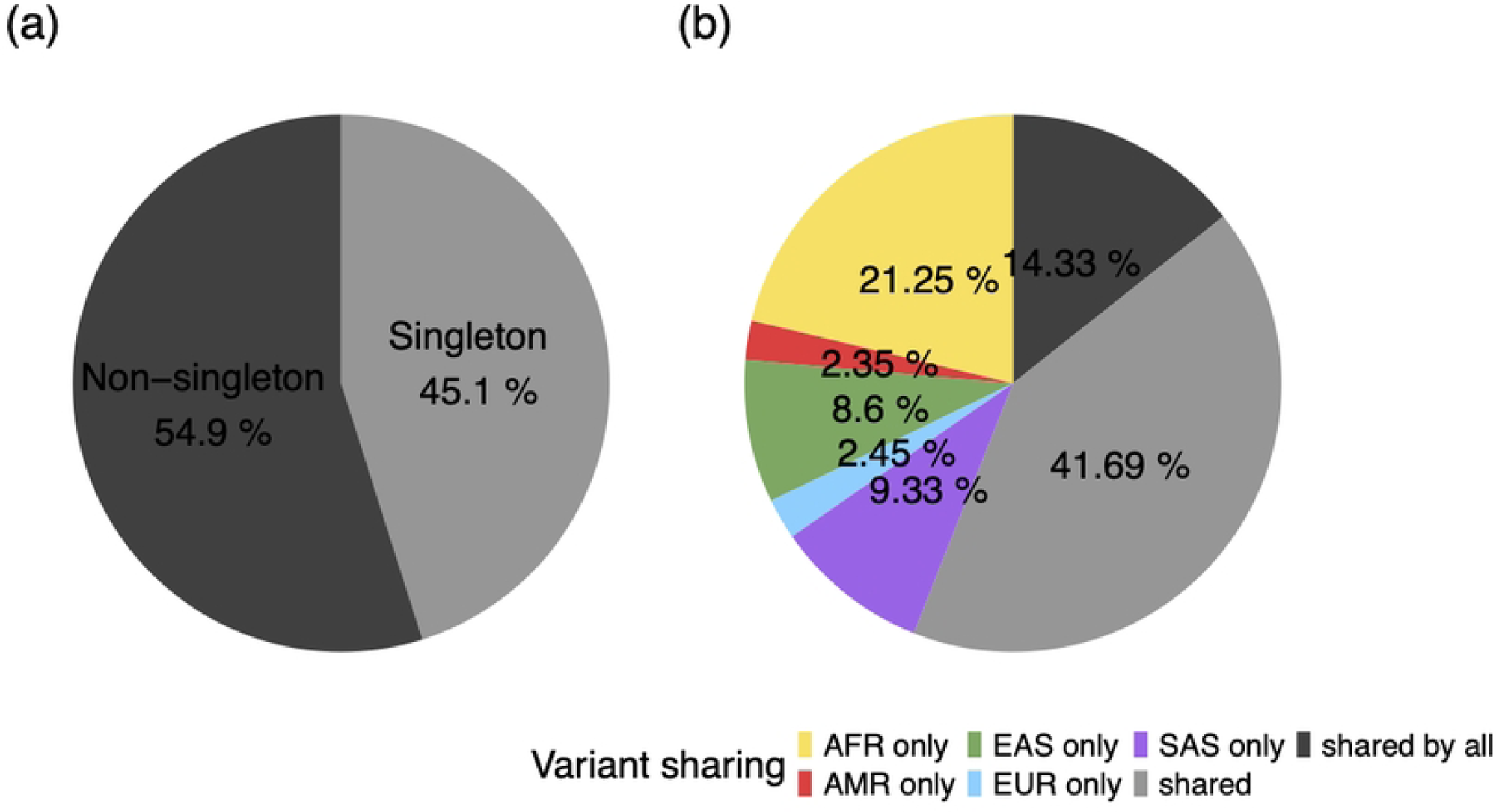
Distribution of biallelic *AHR* SNPs in The 1000 Genomes Project. (a) Majority of *AHR* SNPs (54.9%) occurred in more than one individual (non-singletons); (b) Non-singleton SNPs were mostly shared among 1000 Genomes super-populations (56.02%). AFR: African, AMR: Ad-mixed American, EAS: East Asian, EUR: European, SAS: South Asian super-populations.

### Functional *AHR* variants were shared with the European super-population and frequencies varied widely between super-populations

*AHR* variants with effects on the biological function of AhR have the potential to alter xenobiotic metabolism and are relevant to global public health. For the purposes of this study, we focused on six functional *AHR* variants that were previously linked to GxE by the SWHS and other studies. All six of these risk variants were shared across super-populations (S1 Fig). We also analyzed phased haplotypes – the combination of the six alleles found together on a chromosome for each individual. 32 haplotypes were present within the 1000 Genomes pool. Eight haplotypes appeared in all super-populations, including the four most frequent, and the vast majority of individuals’ haplotypes were shared globally (87.28%) (Fig 2a). We found that 21 haplotypes were shared between Europe and at least one other super-population and these represented 95.99% of the individuals in the 1kGP cohort (Fig 2b). This finding supported our assertion that we can extend the SWHS results to non-European super-populations.

**Fig 2.**
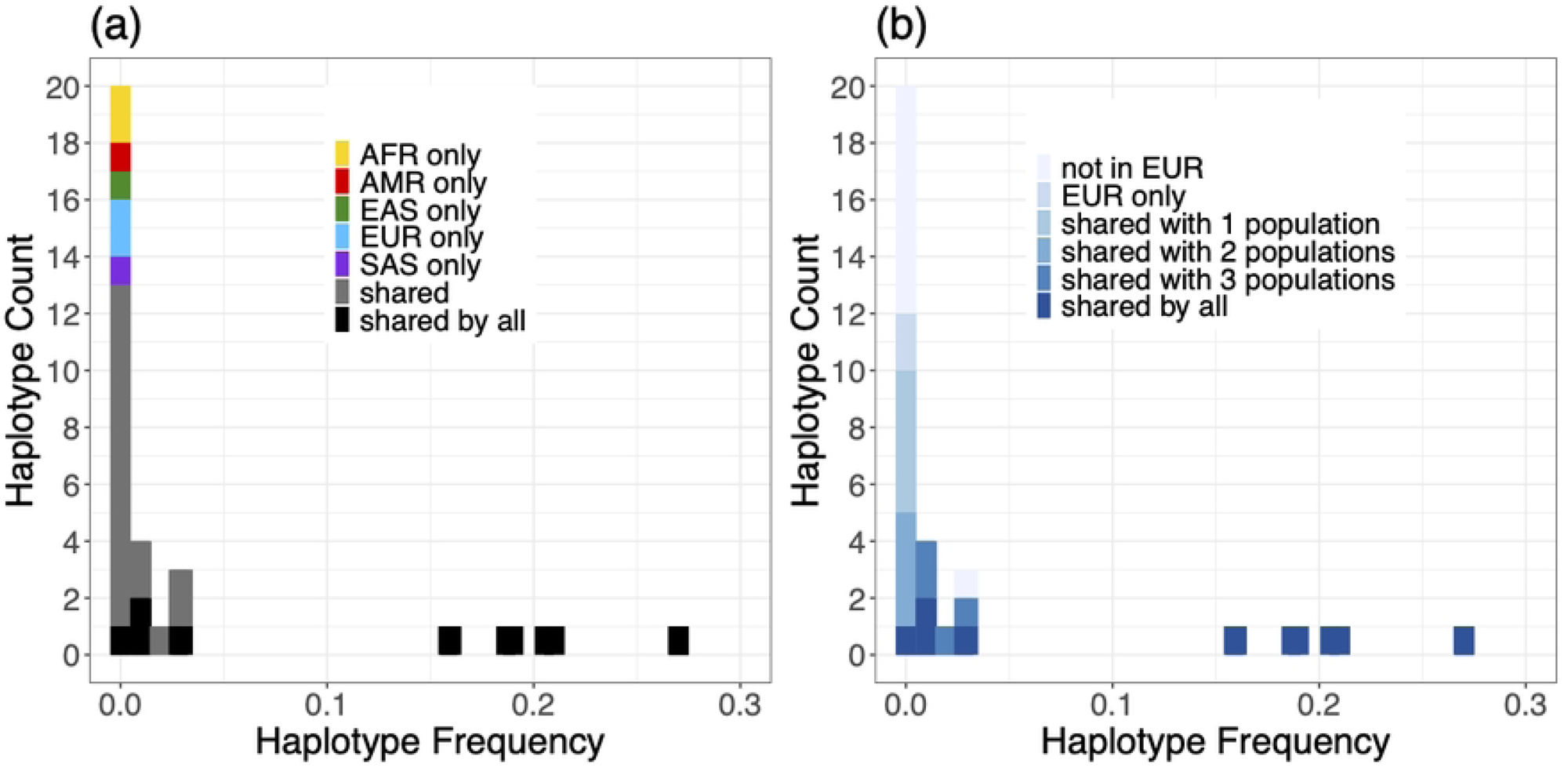
Sharing of thirty-two functional *AHR* haplotypes between 1000 Genomes super-populations. (a) Haplotypes shared between super-populations (99.64%) occurred more frequently than private haplotypes; (b) Twenty-two haplotypes were shared between the European super-population (EUR) and at least one other population. 96% of 1000 Genomes individuals harbored a haplotype that was present in EUR.

The key to our approach is that the risk variant frequencies varied widely between populations (S1 Fig). For example, the risk allele of SNP rs2282885 occurred at a frequency of 98% in African populations, compared to just 60% in European populations. Therefore, we expected differences in population level effects due to GxE. The super-population with the highest and lowest risk variant frequencies varied for each SNP.

### Five of six functional *AHR* variants mediated differential decreases in birthweight among super-populations in simulations

We simulated TCDD-exposed and unexposed populations as described in the Methods. Briefly, we randomly assigned unexposed offspring birthweight from a normal distribution, randomly assigned TCDD exposure level from the distribution reported by the SWHS, and applied fixed GxE effects on birthweight (S1 Table) based on 1kGP genotype and simulated exposure level. Fig 3 shows the spread of mean exposed birthweight from the simulated replicates mediated by each variant. The differences among unexposed populations are purely due to random sampling, since GxE effects were only applied to exposed populations.

**Fig 3.**
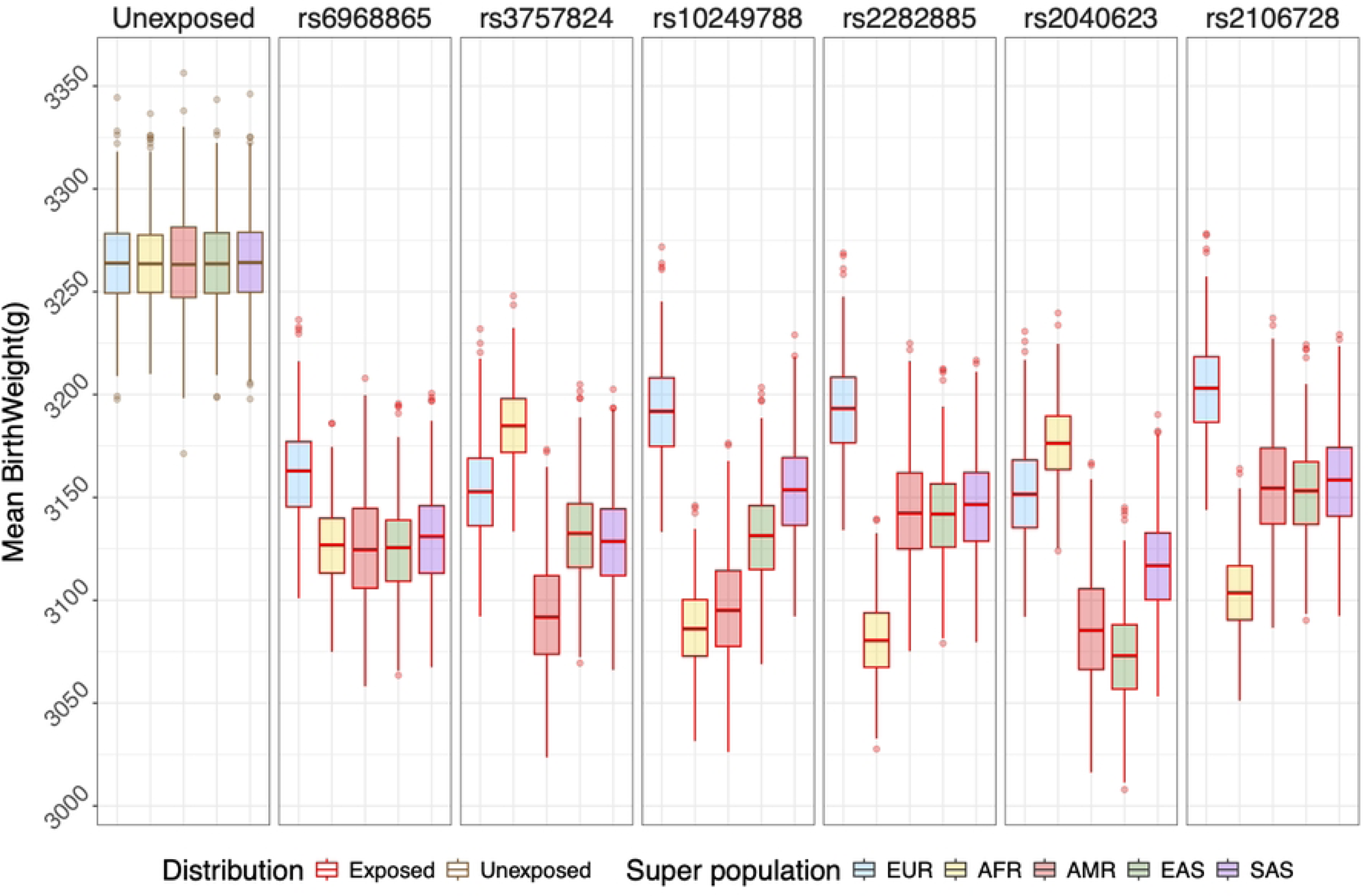
Distributions of birthweight means from simulations of TCDD exposure in 1000 Genomes super-populations. Birthweight means in unexposed populations are shown with a grey border in the first panel (*n*=500 simulated populations per group). Means mediated by GxE involving *AHR* SNPs and simulated TCDD exposure are shown with red borders in the subsequent panels.

We compared randomly selected unexposed and exposed birthweight distributions in 500 t-tests per variant within each super-population. As expected, there were substantial differences in birthweight means between unexposed and exposed populations independent of genetic background. Overall, 92.0% of tests found a significant difference in 15,000 random population pairs (S2b Table). Only five contrasts showed GxE effects in fewer than 90% of tests – rs3757824 in AFR, three SNPs (rs10249788, rs2282885, and rs2106728) in EUR, and rs2106728 in AMR. The first four corresponded to the four lowest risk genotype frequencies (25%-37%) among all the populations. AMR showed a significant effect of rs2106728 only 85.2% of the time, despite an intermediate risk genotype frequency. However, rs2106728 had a smaller effect size than most of the other GxE SNPs, and the AMR population sample was substantially smaller than the four other super-populations. Lower GxE effect size, low risk genotype frequencies, and smaller population sizes could affect the ability to detect the exposure effect in any individual population. Nonetheless, we concluded that in most scenarios we can successfully detect the effect of TCDD exposure in simulated populations after randomly modeling baseline birthweight and exposure level.

Next, we tested if risk genotype frequency differences between super-populations (S3 Table) were sufficient to cause differential birthweight distributions. Briefly, we randomly chose one simulated population from each of the five super-populations and tested for differences in mean birthweight. We sampled without replacement and retested until we had tested 500 times for each of the six SNPs. We found that GxE caused differences among the super-populations for five of the six variants (one-way ANOVA, FDR < 0.05) (Fig 4, S4 Table). We saw no differences in the model for rs6968865, which corresponded to the smallest range of risk genotypes across the populations (0.61 to 0.83). The other variants showed differences in a substantial proportion of tests significant at *p*_adj_≤0.05, ranging from 41.4% to 82.8% of tests (Fig 4, S4 Table). At rs3757824, the AMR super-population had the highest risk genotype frequency (60%), while the AFR super-population had the lowest risk genotype frequency (28%) (S3 Table). That corresponded to where these super-populations fell in terms of TCDD-mediated birthweight decreases with values of - 171.43g and -79.75g respectively (Fig 3, S2a Table). Genotype frequencies for rs2040623 were very similar to those at rs3757824, with the exception that EAS had a higher risk genotype frequency (S1 Fig, S3 Table). We therefore saw a similar pattern at this SNP, but both AMR and EAS showed lower birthweights. At rs10249788, the AFR (-177.69g) and AMR (-168.06g) super-populations and the had the highest mean birthweight decrease and risk genotype frequencies (62% and 58% respectively) relative to the other super-populations. At both rs2282885 and rs2106728, AFR had the highest risk genotype frequencies and the largest effects on the birthweight distribution. We conclude that difference in risk genotype frequencies between populations are sufficient to drive population-level trait differences due to gene-environment interactions.

**Fig 4.**
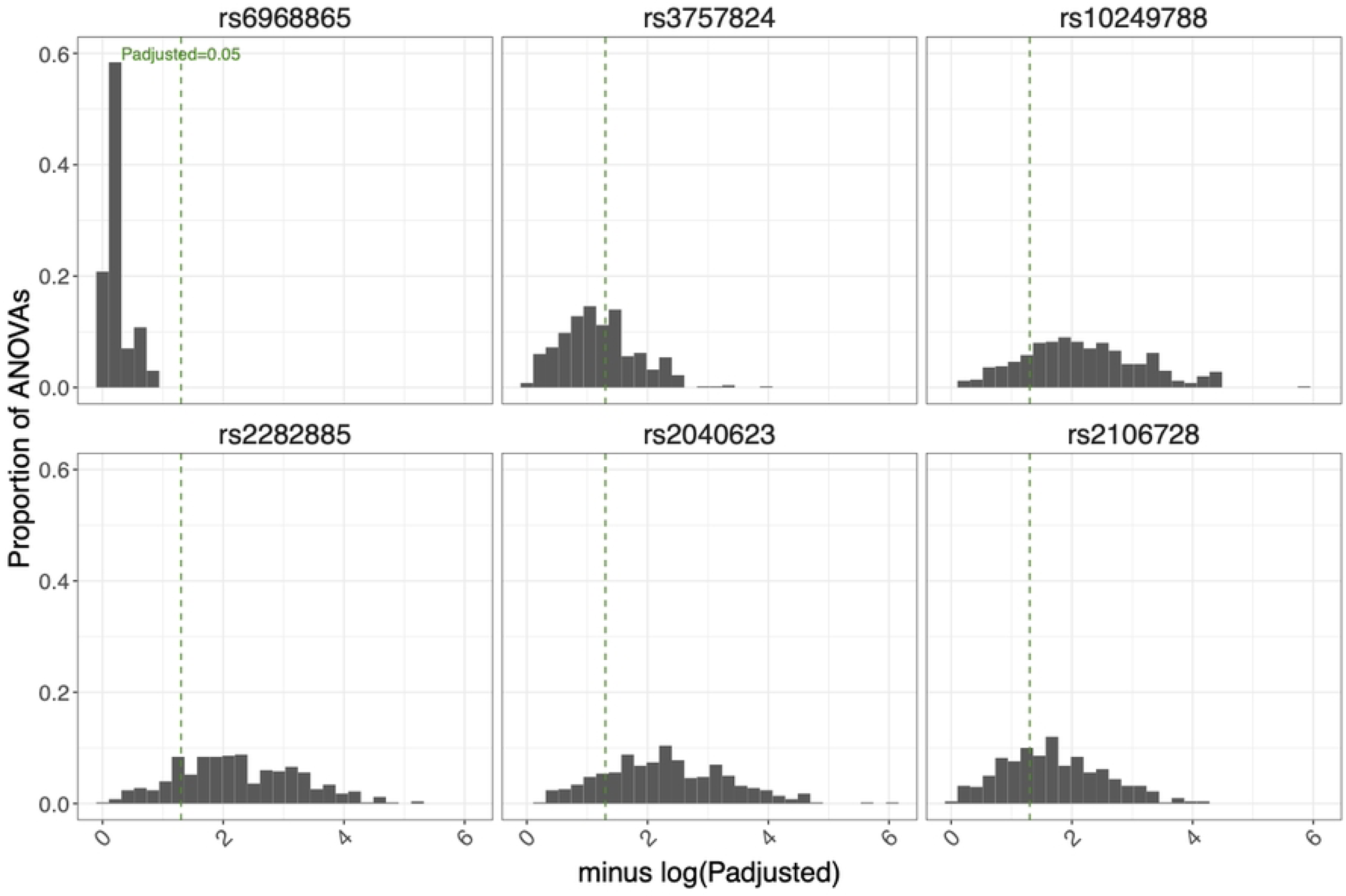
Distributions of FDR-adjusted p-values from comparisons across super-populations. We compared exposed birthweights mediated by *AHR* SNPs and simulated TCDD exposures across 1kGP super-populations. Bins to the right of the p-value threshold (0.05) show statistical tests (ANOVA) where GxE mediated differential decreases in birthweight between simulated populations. Risk genotype frequency differences at five of six SNPs (all except rs6968865) were sufficient to drive differences between super-populations.

## Discussion

We found that most common *AHR* SNPs were shared across global super-populations, including some SNPs that had previously been linked to interactions with specific chemical toxicants that include *AHR* ligands. We built a model to demonstrate how these shared variants could affect the impacts of these exposures on global public health. The importance of AHR function in environmental response, documented GxE linked to *AHR* variants, and the ubiquity of relevant exposures motivated our study. Most known GxE associations in the AHR pathway have been discovered in populations exposed to high doses of toxicants such as the SWHS population, populations exposed to Agent Orange in Vietnam (7, 11, 18), and populations exposed to PCBs and polychlorinated dibenzofurans from contaminated rice bran oil in Japan (19). However, exposures to AHR ligands like PAHs from cigarette smoke and PCBs, and dioxins from dietary intake are common all over the world (20–22). Consequently, the effects of *AHR* risk variants are pertinent to the health of global populations exposed to these common ligands.

The *AHR* gene is well-known for its key role in toxicant metabolism, yet most of the over 13,000 genetic variants found in the gene are unexplored. We calculated two measures of diversity and found that *AHR* nucleotide polymorphism and nucleotide diversity fell within typical ranges for the human genome (P = 2.97×10^-2^, π = 7.5×10^-4^ - 1.25×10^-3^) (4, 23, 24). Sequence variation in other environmentally responsive genes (π = 6.7×10^-4^) and nuclear hormone receptors (π = 4.1×10^-4^) are in the same range (25, 26). In fact, *AHR* looks very much like a typical gene in these respects. The functional consequences of majority of these variants remain unknown.

We illustrated our risk prediction approach on six functional SNPs previously linked to GxE interactions by the well-established SWHS. Our study is the first to estimate population risk due to GxE interactions involving an environmental exposure. A similar study used the same principles to predict variation in Alzheimer’s risk in multiple populations, based on variants identified in genetic association studies (27). An extension of our approach would be to apply the same models to ∼70 *AHR* variants that have been associated with traits in the GWAS catalog (2). A few of these SNPs have also been implicated in GxE, including interactions with coffee consumption and cigarette smoking (28, 29), and with adverse health outcomes ranging from decreased birth size to colorectal cancer risk (12, 30). Our approach could be extended to additional *AHR* candidate SNPs (31–33).

We posited that global super-populations would have disparate outcomes after toxicant exposure if risk variant frequencies differed among those populations. Like so many genetic associations in public data sources, these candidate SNPs were identified in a European cohort. The fundamental assumption of our work is that GxE effects from a population in Seveso, Italy would apply to other populations. This assumption was necessary since that study is one of the very few sources of GxE effect estimates. There was additional support for one of those SNPs (rs2282885) that was associated with differential PAH metabolism in Chinese coke oven workers (10), which also demonstrated how GxE can be shared across populations.

Our simulations were robust to random assignment of birthweight and TCDD exposure doses. In some simulations the most susceptible individuals in the population will be randomly assigned a high birthweight, low exposure, or both. In those cases, the TCDD effect was not detectable, but we found it was detectable the vast majority of the time. Our model predicted that five *AHR* SNPs would result in population-specific birthweight differences due to GxE after TCDD exposure. We had no prior expectation of which super-population would have highest or lowest GxE risk. In fact, the super-population with the most GxE risk varied from SNP to SNP. For example, AFR had the lowest risk genotype frequencies at two SNPs, while having the highest risk genotype frequencies at three others. At the sixth SNP, we found no differences between populations, and this corresponded to the smallest range of risk genotype frequencies. The specific thresholds at which genotype frequency differences map to significant birthweight differences would depend on several factors including the trait variance, GxE effect size, and population size.

Epidemiological studies that incorporate both genetic and environmental factors are rare. Our approach attempts to extend those results to the global 1kGP super-populations. Natural extensions could simulate population-specific effects of *AHR* variation for other relevant exposures such as cigarette smoke or diesel exhaust. Population-level polygenic risk scores based on risk allele frequencies have explained trait variation in the 1kGP (34), and those approaches could possibly be extended to address GxE. Our basic framework could be applied to a wide variety of other toxicants, candidate genes, and health outcomes.

## Conclusions

In conclusion, genetic variation that governs susceptibility to environmental factors is abundant and often shared across global populations. We used differences in risk variant frequencies to show how some populations have higher predicted risk relative to the well-studied European population. While studies in understudied populations provide opportunities to discover novel variants, integrating GxE studies from exposed individuals with global population genetic data can help us elucidate population differences in susceptibility. Consequently, this will inform risk assessment strategies and advance global public health.

## Supporting Information

**S1 Fig. Frequencies of functional *AHR* SNP alleles associated with 2, 3, 7, 8-tetrachlorodibenzo-p-dioxin (TCDD) effect on birthweight across global super-populations**. Risk alleles were shared between super-populations but occurred at different frequencies.

**S2 Fig. Distributions of simulated TCDD concentration in 1000 Genomes super-populations.** Truncated log-normal distributions were simulated to model exposure in SWHS with a median of 55.9 parts-per-trillion (ppt), range of 2.5 to 56000ppt, and each super-population’s sample number.

**S3 Fig. Distributions of FDR-adjusted p-values from comparisons of simulated unexposed and exposed populations.** T-tests compared unexposed and exposed birthweights mediated by *AHR* SNPs within each 1000 Genomes super-population (*n*=500 per group). Bins to the right of the p-value threshold (0.05) show simulations where the GxE effect of the SNP mediated significant decreases in predicted birthweight after simulated TCDD exposure.

**S1 Table. Six maternally functional SNPs in the aryl hydrocarbon receptor gene interacted with TCDD to influence offspring birthweight in SWHS data.** GxE effect was calculated as the difference in SNP effect on birthweight between individuals with the risk and non-risk genotypes.

**S2 Table. Changes in mean birthweight mediated by *AHR* SNPs after simulated TCDD exposure in The 1000 Genomes Project super-populations.** (a) Mean birthweight differences (g) derived from 500 simulations of exposed and unexposed populations for each super-population. (b) Proportions of 500 t-tests significant at a false discovery rate (FDR) threshold of 0.05.

**S3 Table. Genotype Frequencies by 1kGP Super-Population for *AHR* risk SNPs.**

**S4 Table. Counts of FDR-adjusted p-values from comparisons across super-populations.** We simulated exposed super-populations and compared mean birthweight across them using analyses-of variance (ANOVAs) (*n*=500 simulations) Substantial proportions of were below an adjusted p-value threshold of 0.05 for five of six SNPs. P-values were adjusted using the Benjamini-Hochberg false discovery rate method. ANOVAs compared birthweights mediated by *AHR* SNPs after simulated TCDD exposure across 1000 Genomes super-populations.

